# Robust Reconstruction of CRISPR and Tumor Lineage Using Depth Metrics

**DOI:** 10.1101/609107

**Authors:** Ken Sugino, Tzumin Lee

## Abstract

Lineage reconstruction using CRISPR edited barcodes are becoming wide-spread and methods robust against noise are in need. Neighbor-Joining (NJ) algorithm is a robust distance based algorithm extensively used in phylogeny field. NJ is also used for CRISPR-encoded-lineage (CEL) reconstruction with proper re-rooting since NJ is unrooted algorithm. However, we found NJ works without re-rooting for reconstructing CEL when the lineage contains multiple trees but not for a single tree. Examining why this is the case leads to the idea of “depth metrics”. The notion of depth metrics also naturally explains why Russell-Rao metric, previously found best metric for CEL reconstruction, works well. Furthermore, based on the probabilistic model of CEL, we constructed a new metric that performs better than Russell-Rao metric. We also propose inferring ancestral code during reconstruction instead of using a linkage method. These, together with Nearest-Neighbor-Interchange resulted in a new robust method for reconstructing CEL or tumor-cell-lineages which share same assumptions as CEL.

## Introduction

Recent development of CRISPR technologies and single cell sequencing opened the possibility of reconstructing cell lineages using somatic mutations caused by CRISPR edits during development. However, a caveat of single cell sequencing is low sensitivity and this results in loss of information on CRISPR edits. Thus, it is imperative to have an algorithm that can reconstruct accurate lineage trees even when there are noise in inputs caused by substantial amount of missing values.

Neighbor Joining (NJ) [1] is the most widely used distance based tree reconstruction algorithm. It is also known to be robust against noise [2]. NJ is unrooted, so for application to CRISPR-coded lineage reconstruction, unedited (root) node has to be added to the input and the output has to be reordered so that the unedited node becomes the root (re-rooting). Although NJ is unrooted, the output of the algorithm is in a rooted format. We found that for inputs consisting with multiple independent trees, NJ delivers correct outputs without re-rooting. By investigating why this is the case, we found Q-matrix of NJ approximates depth matrix, which consists of the depth of least common ancestors, of the input tree when the input consists of multiple independent trees. Analogously, Russell-Rao metric [3] or SharedEdits metric [4] calculates approximate depth matrix when mutations are “irreversible” and the state of the root node (i.e. unedited for CRISPR-barcodes) is known.

Based on these findings, we created a better bottom-up reconstruction method than the complete linkage method which was found to be best previously [4] by directly assigning inferred code to the new node representing parent of nodes being merged. Further, we calculated probabilistic structure when SharedEdits metric fails to faithfully approximate depth matrix and created an improved metric with less failure in the case of CRISPR-barcodes.

Finally, we implemented Nearest-Neighbor-Interchange (NNI) as a post-bottom-up-reconstruction refinement of tree topology. These resulted in an accurate, fast and robust algorithm suitable for use in CRISPR-coded lineage reconstruction.

## Results

### Why does Neighbor-Joining without re-rooting work for multiple trees?

NJ is an algorithm which produces unrooted tree. For CRISPR encoded lineage reconstruction, we need to recover a rooted tree, root being unedited code. Although NJ is unrooted, the algorithm naturally produces rooted output. Therefore, proper root node must be selected and node relationship (the parent-children relationship) has to be rearranged (re-rooting). When proper root state is known as in the case of CRISPR encoding (un-edited barcode) or tumor lineage (wild type sequence), the root state is included in the input and the output is re-rooted so that the node with the root state becomes the top (root) node. We noticed, however, that NJ outputs are correct even without re-rooting when multiple trees starting from unedited root nodes are included in the input (Fig.1A). To see why this is the case, we plotted Q-transformed matrices (Fig.1B) and noticed that the diagonal location with the highest value, which corresponds to the top (root) node of the output, is different between single tree and multiple trees (Fig.1B).The Q-transformation of distance between node i and j (*d_ij_*) is *d_ij_* subtracted by average distances from i and j to all other nodes (*a_i_* and *a_j_* respectively):

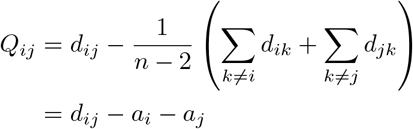

**Figure 1:**
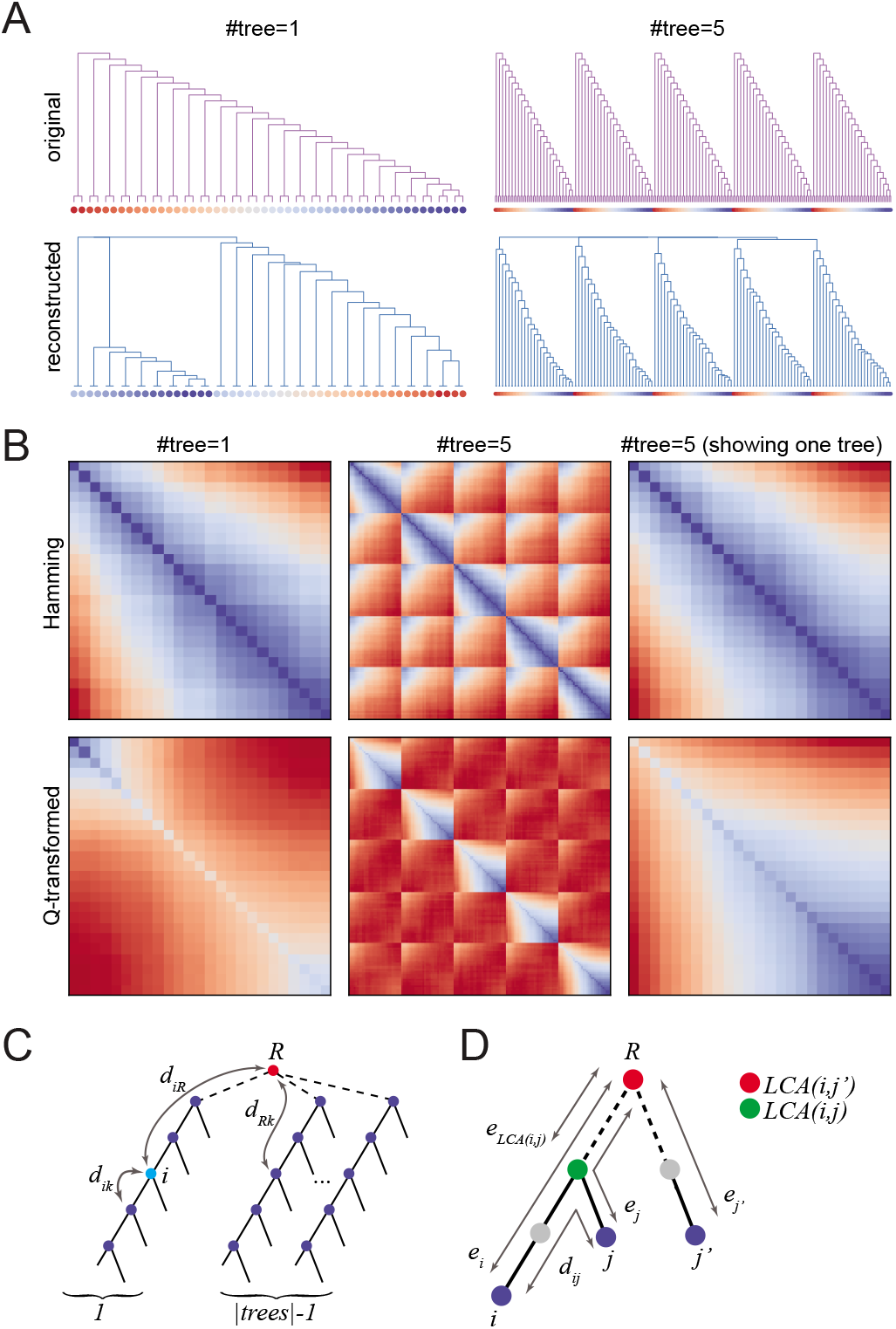
NJ reconstruction without re-rooting. (A) Top: original inputs. Bottom: NJ reconstructed outputs without re-rooting (B) Top: Hamming distances between leaves. Bottom: Q-transformed values between leaves. (C) A cartoon showing the decomposition of *d_ik_* between tree distances. (D) A diagram showing relationship between *d_ij_* − *e_i_* − *e_j_* and −2*e*_*LCA*(*i, j*)_.

(Where *n* is total number of nodes.) In the case of multi-tree case, the average distance from node i to all other nodes (*a_i_*) can be rewritten as the sum of within tree and between tree components. When the number of trees are large, it converges to distance from root (*R*) and node *i* (*d_iR_*) plus some constant since the first term in eq.1 becomes small, the coefficient of the second term (|*k* ∉ *tree*(*i*) | / Σ_*tree*_ |*tree*| − 2) becomes close to one, and the third term which can be rewritten as third and fourth terms in eq.2 (*BL_tree_* = Σ_*k*∈*tree*_ *d_Rk_*) becomes constant since the fourth term becomes small (see Fig.1C). (*tree*(*i*) indicates the subtree node *i* belongs to when the root is removed.)

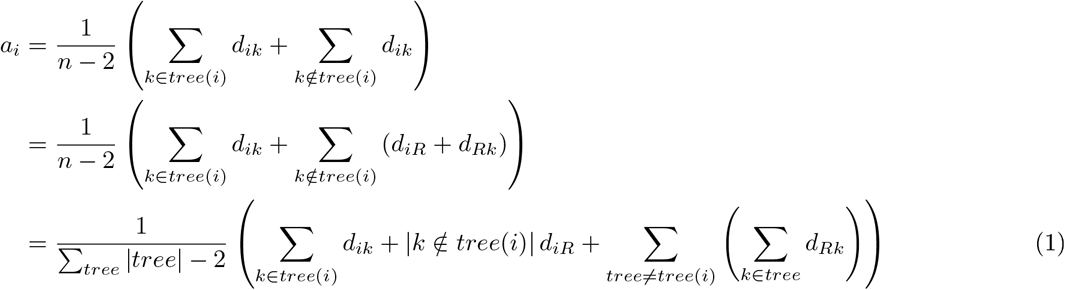

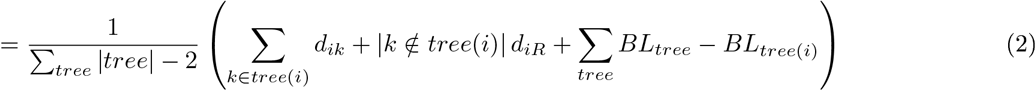

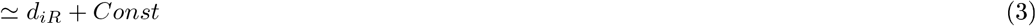

Rewriting *Q_ij_* using this approximation and rewriting *d_iR_* as just *e_i_, Q_ij_* can be expressed as follows (see Fig.1D):

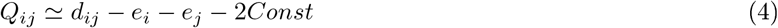

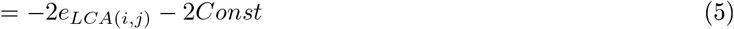
where *LCA*(*i, j*) is the least common ancestor of node *i* and node *j*. This means that the (*i, j*) element of the Q-matrix represents the depth of the least common ancestor of node *i* and *j*. (Deeper depth being smaller.) When *i, j* belongs to different trees, the least common ancestor is the root, so *Q_ij_* is close to −2*Const* and is larger than *Q_ij_* when *i, j* belongs to a same tree, since *Q_ij_* is the depth of the least common ancestor of node *i, j* which is always deeper than the root.

The NJ algorithm reconstructs a tree by iteratively merging the pair with lowest *Q*. With the above approximation, we can interpret this procedure as building a tree from its deepest part. This is why NJ correctly reconstruct rooted tree without re-rooting when input consists of (many) multiple trees. When input is a single tree or small number of multiple trees, this is not the case, so NJ does not output correctly rooted tree. In the next section, we will look in more detail at the quantity *Q_ij_* approximates: the depth of least common ancestor which we call “Depth Matrix”.

### Russell-Rao (SharedEdits) metrics is a Depth metric

In the previous section, we defined “depth matrix” of a tree, whose (*i, j*) element is the “depth” of the least common ancestor of node *i* and node *j*. Depth is a map from nodes to real number: *i* ↦ *d*(*i*) that satisfies *d*(*i*) > *d*(*j*) when node *i* is an ancestor of node *j* (i.e. deeper the smaller). With this notation, depth matrix is a map from pairs of nodes (*i, j*) to real number:

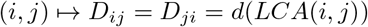

Since *LCA*(*i, i*) = *i*, *D_ii_* = *d*(*i*), that is, diagonal elements of depth matrix are depth themselves. Since *d*(*LCA*(*i, j*)) ≥ *d*(*i*), *d*(*j*), *D_ij_* is always bigger than *D_ii_* and *D_jj_*. Or:

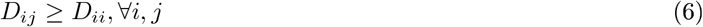

For Type1 (linear) tree, depth matrix has inverted L-shaped pattern when depth of node *i* is defined as negative branch length from root to node *i* (Fig.2A).

**Figure 2:**
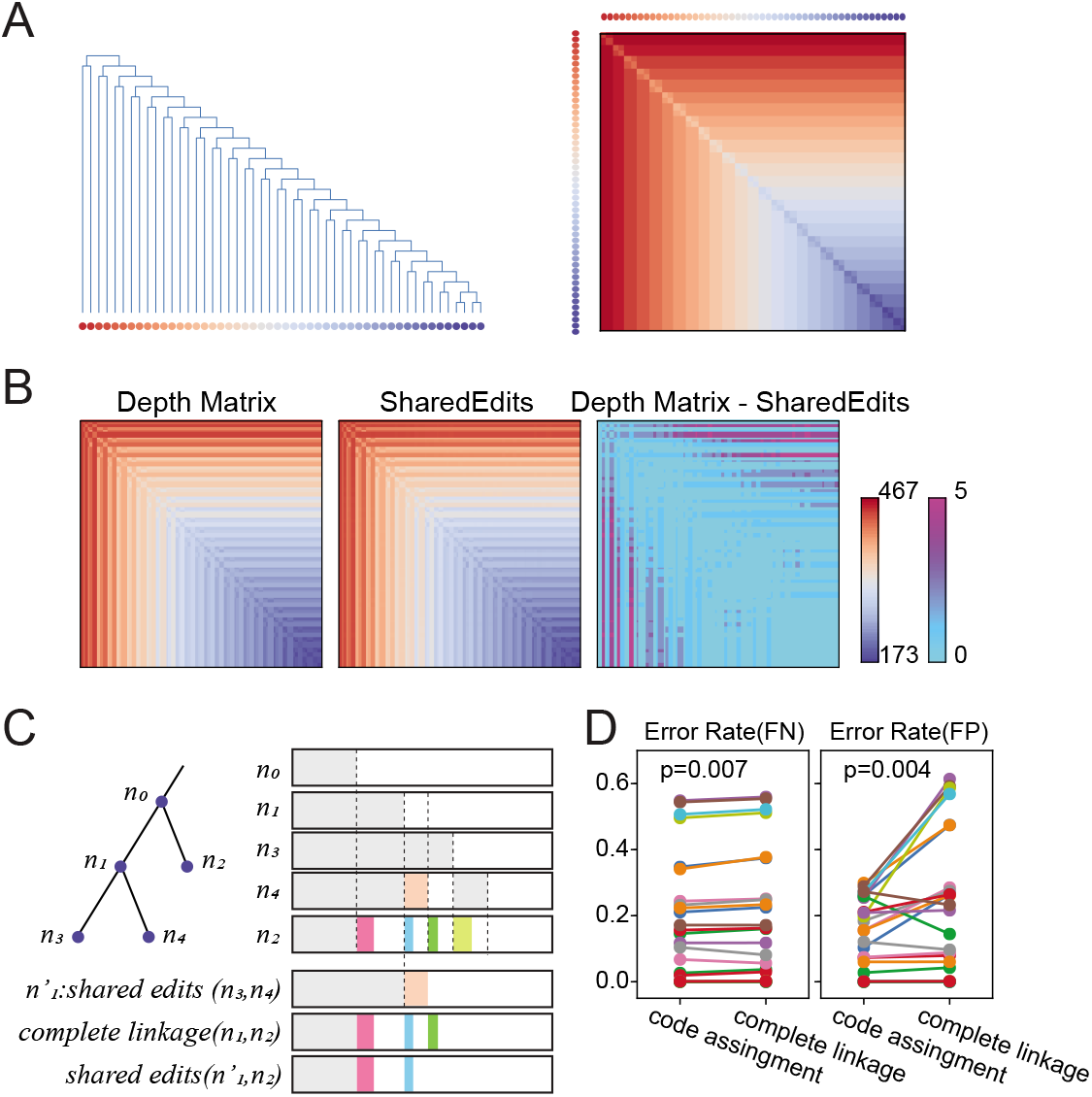
Depth Matrix. (A) (Left) Type1 tree structure. Leaf nodes shown at the bottom are colored according to their depth, shallow (red) to deep (blue). (Right) Depth matrix between leaf nodes using negative path length from the root as depth. Matrix elements are ordered according to depth as shown on the sides. Color indicates larger (red) to smaller (blue). (B) (Left) Depth matrix calculated using number of edits as the depth. The tree is Type1 tree of length 25 with number of units 467 and edit rate of 0.04 (which guarantees some edits at every step more than 99.9% of the time. Unlike (A) values for all nodes of the tree are shown. (Still ordered by depth). (Middle) Pairwise values calculated by SharedEdits metric. (Right) Difference between the depth matrix and SharedEdits matrix. (C) Due to coincident edits (colored portion), ancestor edit number is overestimated by SharedEdits metric (e.g. *n*_1_ edit number is overestimated by orange part). This can be seen in (B) right panel as well (0 ~ 5 overestimation). When calculating metric between merged nodes (*n*_1_) and other node (*n*_2_), assigning shared edits of *n*_3_ and *n*_4_ to *n*_1_ (denoted as 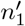) can reduce the overestimation compared to complete linkage (which takes minimum SharedEdits of members). (D) Performances of code assignment and complete linkage are compared using the benchmark datasets used in [4]. Code assignment performs significantly better (paired t-test). FN: False Negative, FP: False Positive.

When we have a depth matrix of a tree, we can reconstruct the original tree from the matrix. First, we observe that a pair of leaf nodes (*i, j*), *i* ≠ *j* with the smallest *D_ij_* is a cherry in the tree, since if not, then there should be another node *k* between *LCA*(*i, j*) and either *i* or *j* and *D_ik_* or *D_jk_* should be smaller than *D_ij_*. So the pair with smallest *D_ij_* should be merged to create a new internal node *k* and its depth should be *d*(*k*) = *D_ij_*, since this new node is the least common ancestor (parent) of nodes *i, j*. Now from the depth matrix, we delete two rows and columns corresponding to nodes *i, j* and add a new row and a column corresponding to the new parent node *k* of node *i, j*. The new depth matrix elements between node *k* and other nodes are defined as:

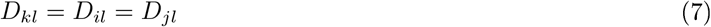
since least common ancestor of node *k* to any other nodes *l* should be same as the least common ancestor between node *l* and nodes *i, j*. This new matrix is a depth matrix of the tree where node *i, j* are replaced with node *k*, which has one less number of nodes. This procedure can be repeated until the number of nodes in the tree becomes one.

We now call any metric of characters that (approximately) reproduces depth matrix of a tree a *depth metric*. Russell-Rao metric (or SharedEdits metric as we expanded the definition to non-binary case and renamed in [4]) calculate one minus the fraction of common characters, or in the case of CRISPR-encoding, one minus the fraction of shred edits. We note that the number of remaining units/targets (number of total units minus number of edited units) is a depth of the lineage tree if every cell cycle (tree branch) has at least one edit, as this number reduces along the depth. The shared edits of cell *i* and *j* is the edits of the least common ancestor of cells *i, j* except for those coincident edits acquired by cell *i* and *j* after their least common ancestor, which are infrequent by chance. Therefore, SharedEdits metric is a depth metric (see Fig.2B).

When reconstructing a tree using a metric in a bottom-up hierarchical manner, it is usually necessary to specify a “linkage method”, which specifies how to calculate metrics between merged node and other nodes from the values of leave nodes. Previously, we found complete linkage method, which takes the maximum of element-wise distances between clusters, performs the best. This corresponds to finding a leaf node with minimal shared edits in the merged cluster. We find, however, that this can still overestimate the true values in some cases (see Fig.2C). To overcome this, instead of using linkage method, we assign the shared edits of the nodes being merged as the code of the newly created node (Fig.2C) and directly calculate SharedEdit distance between the new node and other nodes. When benchmarked against the set of input trees used in the previous study [4], we found ancestral code assignment resulted in better error rates compared to complete linkage method (Fig.2D).

In summary, we describe a notion of depth matrix, and show SharedEdits metrics works well for tree reconstruction because it closely approximates depth matrix. By examining linkage process, we found assigning shared edits code to ancestor improves the reconstruction accuracy.

### Improved SharedEdits metric

SharedEdits metric approximates the depth matrix pretty well (Fig.2B left, middle panels), however there are occasional underestimation of depth due to coincident edits (Fig.2B right panel). This can sometimes cause wrong order of reconstruction and result in error. To understand the nature of these “flips”, we calculated probability of swaps and ties between most closely related two pairs of nodes (3,4 vs. 2,4 in Fig.3A). Since the edits of these nodes are inter-related, and we also need assume there is at least one edit from node 0 to node 1 (*e*_01_ > 0), we need to integrate multinomial distribution of 5 variables (see Fig.3A right panel). We excluded *e*_01_ = 0 cases because no metric can correctly assign depth in these cases and so these cases are irrelevant to metric performance. The resulting swaps and ties probabilities for SharedEdits metric (Fig.3B top row) have peaks when expected number of edits is a little less than 2 indicating swaps and ties are more likely to happen when fewer new edits are possible. However, the peak height is proportional to rate, likely because smaller edit rate yields smaller variability. We also observe higher number of edit outcomes (*L*) results in smaller swap/tie probabilities, likely because larger *L* results in less coincident edits. We also noticed that most of the swaps and ties are actually ties. For example, in the case of *r* = 0.1, *L* = 3, *nU* = 25, *P*(*swap*) = 0.00055 and *P*(*tie*) = 0.015, that is 96% is tie.

**Figure 3:**
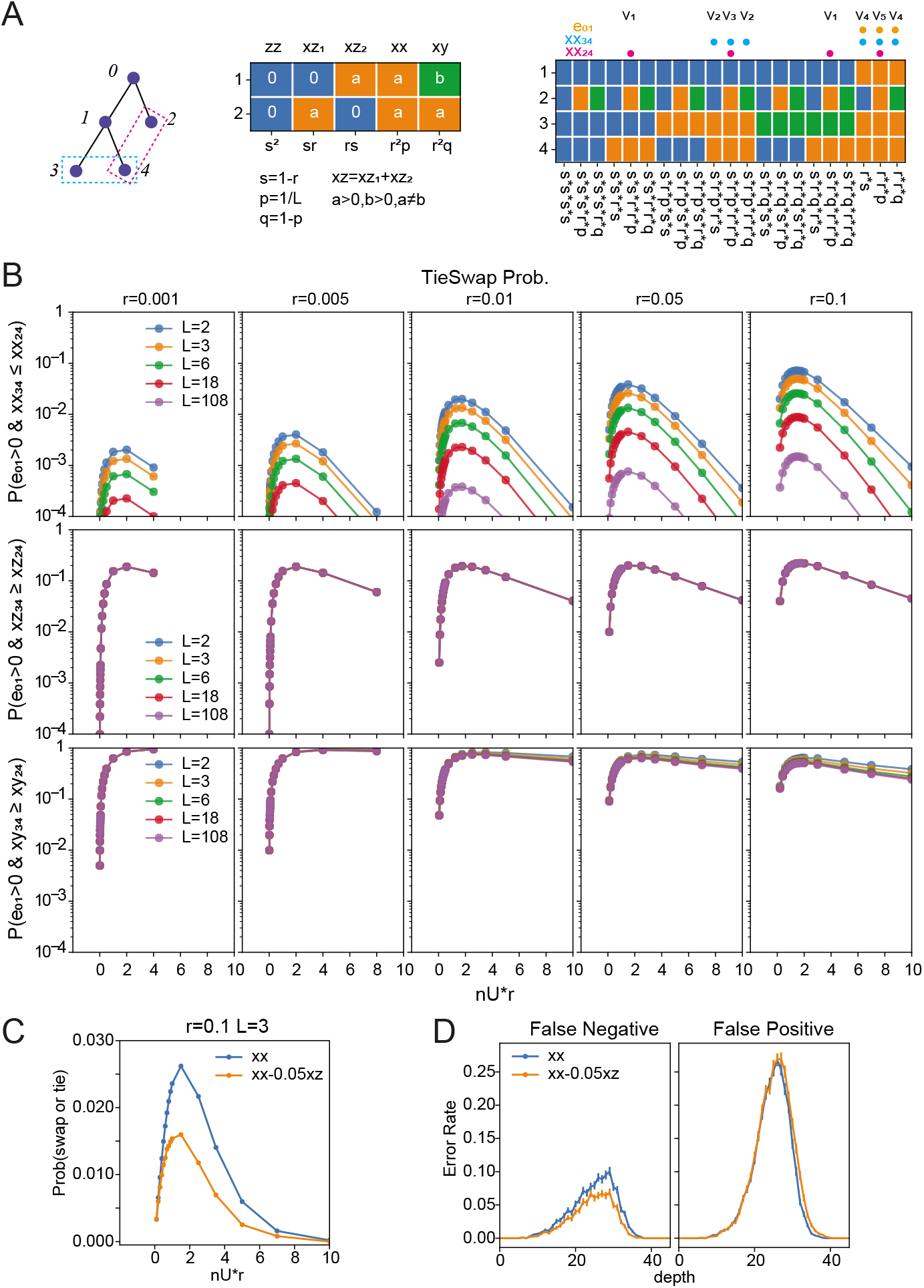
Swap/Tie Probability. (A) (Left) At node 0 there are *nU* unedited units that are edited at the rate of *r* with *L* outcomes. We compare codes for pairs (3, 4) and (2, 4). (Middle) There are four types of cases when two codes are compared: number of units where a) both are unedited (zero) (*zz*), b) one is edited and the other is unedited (*xz*), c) both edited and same (xx), and d) both edited and different (*xy*). These four variables obey multinomial distribution with the rates shown below each column. (Right) When metrics of two pairs (34 and 24) are compared, interdependence has to be taken into account. Random variables *xx*_34_, *xx*_24_, *e*_01_ can be expressed as the sum of other variables (indicated by *v_i_* above columns) which obeys multinomial distribution with the rates shown below the columns. (*e*_01_ is the number of edits from node 0 to node 1.) (B) (Top row) Probability of *xx*_34_ and *xx*_24_ being swapped or tied when there is some edit between node 0 and 1(*e*_01_ > 0) are shown for various combination of edit rates (*r*; columns), edit outcomes (*L*; colors) and number of unedited units at node 0 (*nU*; x-axis but shown as *nU* * *r*, expected number of edits). (Middle, Bottom rows) Similar to top row but for *xz*_34_ ≥ *xz*_24_ and *xy*_34_ ≥ *xy*_24_. (C) Similar to (B) but comparing metrics *xx* and *xx* − 0.05*xz* for the case of *r* = 0.1, *L* = 3. (D) 2000 Type 1 trees with *r* = 0.1, *L* = 3, *nU* = 180 were simulated and nodes with no-edits are removed. Then they were reconstructed using either *xx* (SharedEdits) or *xx* − 0.05*xz* (SharedEdits w/xz), and their false negative and false positive errors are shown.

To see whether other portions of the pair comparison (*xz, xy* in Fig.3A middle) can help in resolving the ties of SharedEdits (*xx*), we first examined how the size of other parameters reflect correct depth relationship (Fig.3B middle/bottom rows). The flip probabilities of *xz* does not depend on *r* nor *L*, and peak height is around 0.2, whereas those of *xy* were much higher. From these, we inferred *xz* may help in resolving some of the ties of *xx*. In fact, the swaps/ties probabilities of random variable *xx* − 0.05*xz* is smaller than that of *xx* (Fig.3C). Moreover, this metric also performed about 30% better than SharedEdits in actual reconstruction (Fig.3D).

In summary, we can improve the performance of the SharedEdits metric by incorporating other components of the pairwise comparison.

### Robust reconstruction

Because of the sensitivity issue of single cell assays, imperfect recovery of codes is a common problem of CRISPR-encoded lineage tracing experiments or tumor lineage tracing. Therefore, it is imperative that any practical reconstruction algorithm is robust against missing codes. To make the current algorithm more robust, we implemented two additional steps. First, during the code assignment, we incorporate code inference when there are missing values (Fig.4A). When one unit is edited and the other is missing (case 7,8 in Fig.4A), we adapted the edited level as the level for merged node. Likelihood of this being correct depends on how many shared edits nodes *i, j* have. It is more likely to have missing code when merging leaf nodes, which are likely to be deep in the tree, so they are usually more likely to have many shared edits. Moreover, even if this assignment is wrong (that is true values is 0; unedited), it is likely to be corrected at the next merging step. Alternatively, it may be useful to track the frequency of the levels being merged as in the case of FastTree [5].

**Figure 4:**
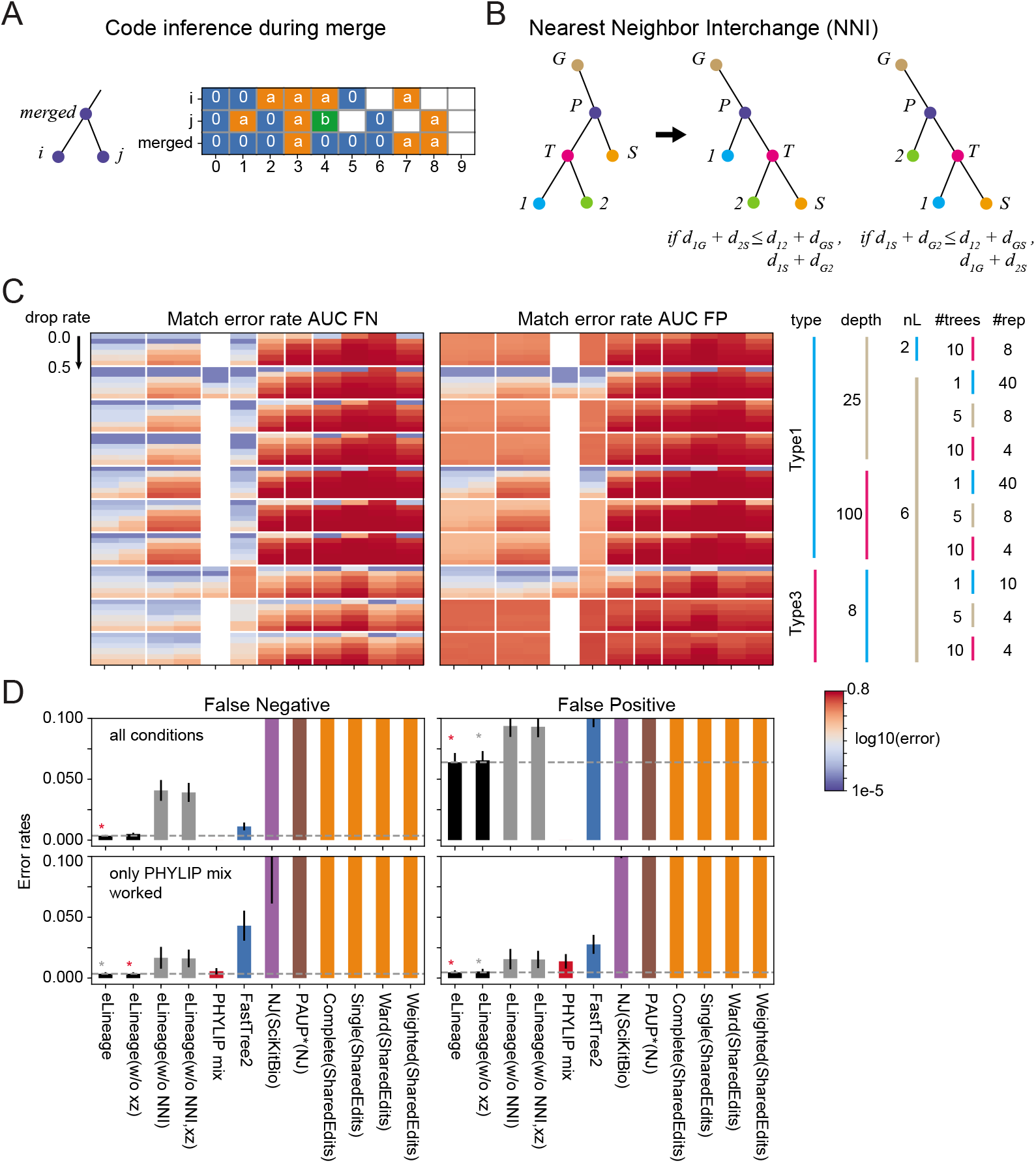
Robust reconstruction against code drops. (A) Code for merged node is inferred from child nodes. 0: unedited, a: edited, b: edited and different from a, white: missing. (B) Nearest neighbor interchange rearranges part of the tree nodes so that total distance of that part becomes minimal. (C) (Top left) mean false positive error rates shown as heatmap in log scale. Y-direction indicates input conditions, which are indicated on the far right. Within each block of condition, code drop rate is gradually changed (from 0 to 0.5 with an increment of 0.1). X-axis legend is shared with (D). White indicates that program did not finish within the upper time limit of 20 min. (Top right) Same as left for false positive error rates. (D) (top row) Mean error rates are shown for all software packages. PHYLIP mix is not included since it did not finish for some of the conditions. Error bars indicate standard error. Orange star indicate one with minimum error rate. Gray star indicate ones within standard error from the minimum. (bottom row) Same as top row, except only conditions where PHYLIP mix is successfully completed are used.

In addition to code inference, we also implemented Nearest Neighbor Interchange (NNI) as the post-reconstruction correction (Fig.4B). For every internal node *T*, we rearrange its child nodes (1, 2) and sibling node (*S*) so that total distances within the subtree consisting of parent (*P*) and grand parent (*G*) becomes minimal (Fig.4B).

To see the performance of the resulting algorithm, we benchmarked several software packages against various input types with varying degree of code drops (from 0.1 ~ 0.5) (Fig.4C). Overall, the algorithm in the current study, which we named “eLineage”, performs the best. The *xz* correction introduced in the previous section helps in some cases such as Type1 depth 100 with high dropout rates. The NNI greatly improves error rate for the cases with code dropouts. The PHYLIP mix (which implements maximum parsimony algorithm) performs 2nd best. However, the execution time is more than 100 fold than other programs (Fig.Supp1A,B) and in most of the cases where the numbers of leaves are large, it could not finish in the required upper limit of 20 min.

## Discussion

### An interpretation of Neighbor Joining Q-matrix

Neighbor-Joining [1] is arguably the most used distance based tree reconstruction algorithm in phylogeny field. However, the interpretation of what the Q-transformation does has not been straight-forward [6]. NJ is an algorithm that produces unrooted tree. Thus, for the application to CRISPR-code lineages, one needs to include the unedited (root) node in the input and then re-root the output tree so that the included unedited node becomes the top root node. However, we found that this re-rooting is not necessary for cases where many independent trees are included in the input. By asking why this is the case, we found that NJ for multiple trees calculates a depth matrix, (i,j) element of which is the depth of the least common ancestor of node i and j, depth being measured from the unedited root node. For single tree case, we can interpret Q-transform to calculate depth matrix from the node which represents the average of all nodes.

The formula (4) *Q_ij_* ≃ *d_ij_* − *d_iR_* − *d_jR_* + *C* gives rise to true depth matrix when *d_ij_* are true path length between node i and j. However, usually Hamming distance are used instead. In this case, using notation of Fig.3A middle panel, *Q_ij_* ≃ *C* + (*xz*_1_ + *xz*_2_ + *xy*) − (*xz*_1_ + *xx* + *xy*) − (*xz*_2_ + *xx* + *xy*) = *C* − 2(*xx* + 0.5*xy*). That is, Q with hamming distance has a correction term, 0.5*xy*, to the SharedEdits *xx* (when distance is normalized). The positive coefficient is the correct direction considering the probabilistic behavior of *xy* swap/ties (Fig.3B bottom row). We have evaluated this metric and found that it tends to fail to distinguish multiple trees, likely because the value of 0.5*xy* is often larger than *xx* at the top of each subtree (where no code inheritance, which contributes to *xx*, exists) and subverts the distance relationship determined by *xx*.

### Russell-Rao (SharedEdits) metric are Depth metric

It is quite easy to understand that if one has a depth matrix of a tree or close approximation of it, then it is possible to reconstruct the tree accurately. The Russell-Rao metric or its expansion to non-binary case (SharedEdits) calculate number of commonly shared edits between two nodes. These shared edits are inherited from the least common ancestor of the two nodes except for the coincident edits. The number of total edits are depth if there are at least one edits between nodes. Therefore, Russell-Rao or SharedEdits metrics calculate depth matrix except for the factor contributed by coincident edits.

To evaluate the error in reconstruction caused by coincident edits, we calculated probability where depth is swapped or tied (Fig.3B top rows) for closest pairs with wide range of editing rate, number of units and number of edit outcomes. In general the swap/tie probabilities are very small (< 0.1, Fig.3B top) peaking when the average number of new edits is a little less than 2, likely representing the case when there is one coincident edits between node 1 and 2 (in Fig.3A), and another between node 4 and 2.

### Code assignment is better than complete linkage

Previously we found complete linkage works best for Russell-Rao or SharedEdits metric for tree reconstruction [4]. By knowing which value we want to estimate (the total edit number of least common ancestor), we realized that directly assigning shared edits to merged node reduces the effect of coincident edits compared to complete linkage (Fig.2C). This in fact improved the reconstruction accuracy (Fig.2D).

### Improving SharedEdits metric by breaking tie

While calculating the probabilistic behavior of *xx* variable (Fig.3B top), we also found that the swap/tie events were mostly tie events. In the case of *r* = 0.1, *L* = 3, *nU* = 25, 96% of events were tie. This is also consistent with the previous finding [4]. To see whether other variables *xz, xy* can break the tie, we first calculated the probability that their size relationship is consistent with depth relationship (i.e. whether *xz*34 is smaller than *xz*24 in Fig.3A left panel, etc.). The results indicated that *xz* may be more useful since it has less swap/tie probability than *xy* or −*xy* To see *xz* can break the ties, we evaluated the swap/tie probabilities of *xx* + 0.05*xz* (Fig.3C). This was 11 variable multinomial integration (which took 1 hour for *r* = 0.1, *L* = 3, *n* = 1, 2,…, 70, 100, 200, 400, 800) and so we did not exhaustively check for other rates and L’s. The coefficient 0.05 is to make the effect of *xz* only show up when *xx* is tied. Expected value of *xz* is *nr*(1 − *r*) ~ *nr* when *r* is small and so when the average expected number of new edits are less than 20, 0.05*xz* would not change the depth relationship determined by *xx*.

The improvement by adding this tie-breaking term was subtle (~30%, Fig.3D) but it was also observable in the case of reconstructing tree from codes with dropouts (Fig.4C).

### Robust tree reconstruction algorithm

Inadequate sensitivity is a common caveat of single cell experiments. This results in part of the code having missing values. To cope with this, we implemented code inference in the code assignment when two nodes are merged (Fig.4A). This resulted in better performance against code drops compared to (standard) NJ or hierarchical clustering (Fig.4C,D, eLineage(w/o NNI)), however, FastTree2, which implements topological rearrangements based on maximum likelihood and Nearest Neighbor Interchange (NNI) after Neighbor-Joining phase performed even better (Fig.4C,D, FastTree2). We, therefore, implemented NNI as a second phase of reconstruction (Fig.4B). This resulted in further improvement in accuracy in the presence of code drops and resulting algorithm (eLineage) out-performed all other algorithms (Fig.4C,D, eLineage).

### Applicability to tumor lineage tracing

Although lineage reconstruction of tumor cells utilizes spontaneous mutations of genomic DNA as in the cases for molecular evolutionary phylogeny, the relatively short duration of tumor development and availability of numerous mutation sites (so called “infinite site assumption”) allow us to assume that these mutations are fixed, i.e. does not get overwritten. This makes the behavior of tumor mutation set very similar to CRISPR-edited codes. Therefore, although we did not test in this study, we believe the present algorithm is equally useful for tumor lineage tracing.

### Further improvement

The speed of the software in this study (evaluated against the datasets used here) is faster than FastTree2 or NJ, PAUP*, PHYLIP mix but slower than scipy hierarchical clustering (Fig.Supp1). However, current software is code in Python scripting language (vs. C in scipy hierarchical clustering method). Therefore, re-coding the algorithm in C/C++ should further improve the speed.

The software scales as 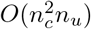 (*n_c_*: number of cells, *n_u_*: number of coding units) and would be slower than FastTree2 for large datasets, which FastTree2 is specifically designed to address. We plan to address this in a future version of the software.

## Materials and Methods

### Error metrics

Error metrics used to evaluate reconstructed trees are as described in previous study [4]. It is a depth dependent expansion of the Robinson-Foulds metric[7]. Briefly, for each depth of ground truth tree, the fraction of the nodes without a matching node in the reconstructed tree is regarded as the false negative error rate. Nodes are matched when the set of leaves under the nodes match. To calculate false positive error rate, ground truth tree and reconstructed tree are swapped.

### Integration of multinomial distributions

Integration of multinomial distribution is done by iterative dynamic programming. It starts from a range that spans the highest probability point (i.e. mean point of the multinomial distribution) and repeatedly expand the region until substantial portion (>99%) of the probability density is covered or integrant (such as *xx*_34_ < *xx*_24_) is deemed to be zero after enough high probability region is covered. Since the number of variables involved in the calculation for even the simplest tree is large (e.g. 20 variables for tree of depth 2, Fig.3A right panel), we created a program to automatically calculate dependencies between variables and associated rates for multinomial distribution. These programs are included in the eLineage software package.

### Simulated datasets used for robustness evaluation

Two types of tree topologies are used. Short (25) and long (100) type1 (linear, asymmetric division, see [4]) and small size (depth 8) type3 (exponential, symmetric division) are tried with varying number of independent trees (1,5 and 10) and edit outcomes (nL; 2 and 6, 2 only for depth 25 type1). Optimum editing rates and number of total units for simple encoding process described in [4] are used. Specifically, for depth 25 tree, *r* = 1/25 = 0.04 and *nU* = 467, for depth 100 tree, *r* = 1/100 = 0.01 and *nU* = 1894, and for depth 8 tree, *r* = 1/8 = 0.125 and *nU* = 143. These rates and number of units guarantees there is at least one new edit 99.9% of the time up to the desired depth. The codes for leaf nodes are used for inputs to the reconstruction algorithms. To evaluate the effect of code dropouts, various portions (10,20,30,40,50%) of the codes are randomly replaced with a value representing missing value (float NaN for eLineage).

### Software packages evaluated

FastTree2 [5], PHYLIP mix [8], PAUP* NJ and UPGMA [9], NJ in scikit-bio (http://scikit-bio.org), hierarchical clustering (linkage) in scipy (http://www.scipy.org) are evaluated. For details of the execution of each package see [4]. The missing values are encoded to adapt to each case (e.g. a missing value is represented as a gap, for fasttree). The eLineage software package include a module which runs each of these programs with appropriate input/output conversions.

### Program availability

The software library implementing the algorithm presented in the current study (eLineage) is available at Bit-Bucket (https://bitbucket.org/kensugino/elineage/). This library also includes scripts to generate tree and simulate CRISPR code editing. It also includes the program used to integrate multinomial distributions described in Fig.3 and runner for other software packages evaluated.

## Supplementary Materials

**Figure Supp.1:**
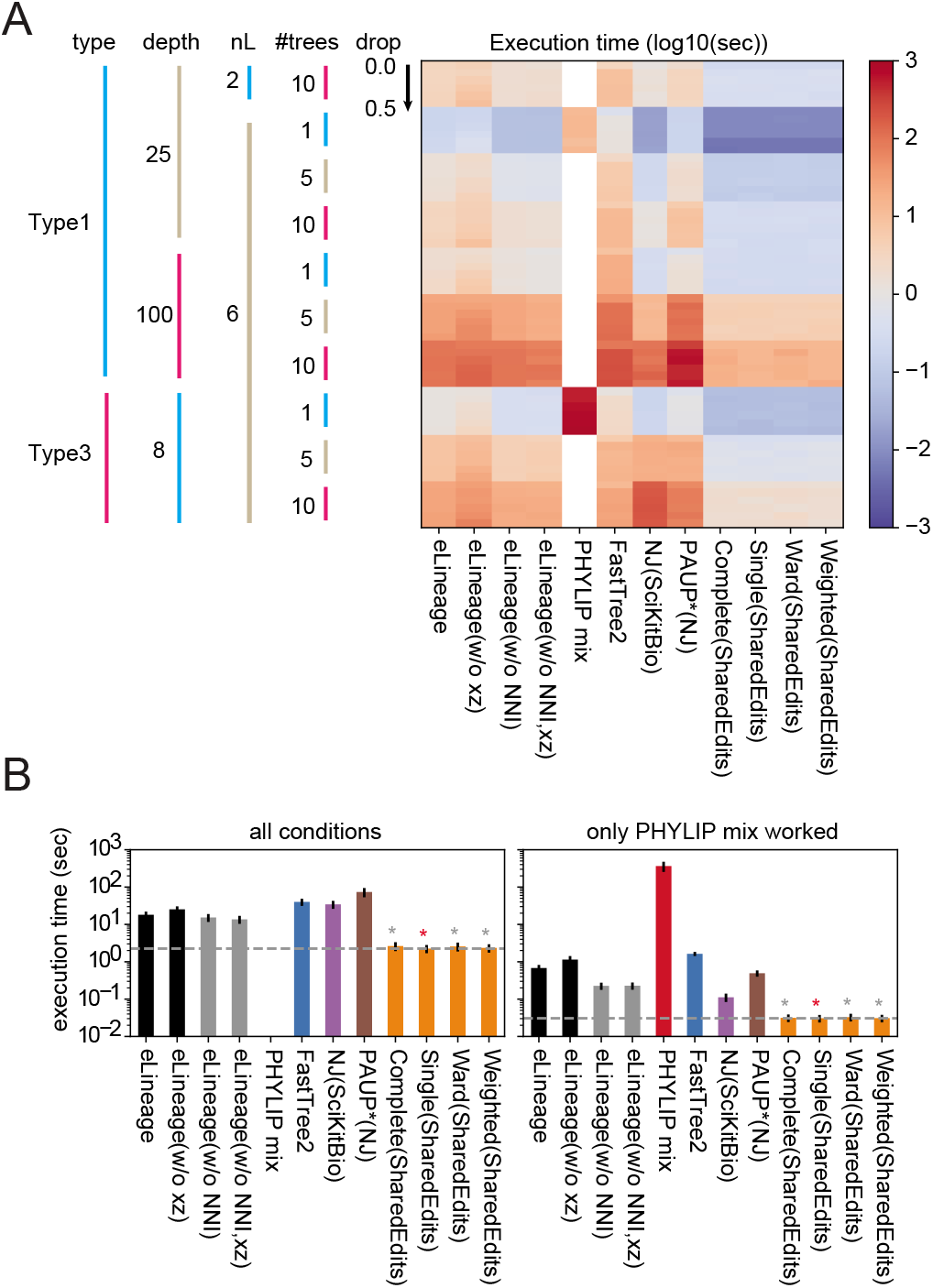
Execution time summary. (A) Heatmap showing execution time (in *log*_10_(*sec*)) for all input conditions and software packages. (B) Mean execution time calculated for all conditions (left) and for only these conditions where PHYLIP mix is successfully completed.

